# A neural code for spatiotemporal context

**DOI:** 10.1101/2022.05.10.491339

**Authors:** Daniel R. Schonhaut, Zahra M. Aghajan, Michael J. Kahana, Itzhak Fried

## Abstract

Time and space are principle organizing dimensions of human experience. Whereas separate lines of study have identified neural correlates of time and space, little is known about how these representations converge during self-guided experience. Here we asked how neurons in the human brain represent time and space concurrently. Subjects fitted with intracranial microelectrodes played a timed navigation game where they alternated between searching for and retrieving objects in a virtual environment. Significant proportions of both time- and place-selective neurons were present during navigation, and distinct time-selective neurons appeared during task-free delays absent movement. We find that temporal and spatial codes are dissociable, with time cells remapping between search and retrieval tasks while place cells maintained stable firing fields. Other neurons tracked the context unique to each task phase, independent of time or space. Together these neuronal classes comprise a biological basis for the cognitive map of spatiotemporal context.

---

Time and space are central organizers of human experience, allowing us to reconstruct the past and envision the future. Lesions to the medial temporal lobe (MTL) and prefrontal cortex (PFC) disrupt associations between events and their temporal (*1, 2*) and spatial (*3–5*) contexts. Parallel lines of research have uncovered neurons in these regions that fire at specific locations within a given environment (‘place cells’ (*6*)) or specific moments within a stable interval (‘time cells’ (*7–9*)), providing a candidate biological basis for the cognitive map of time and space that frames human experience. However, most of our knowledge of these neurons comes from studying them in isolation: place cells are usually recorded during exploratory or goal-directed navigation absent time constraints, while time cells are recorded at fixed locations under timed conditions (*9*). Thus, despite the longstanding assumption that events are bound to combined spatiotemporal contexts in which they occur (*10–12*), it remains unclear how neuronal representations of time and space converge in practice. In addition, although there is now good evidence for place cells in humans (*13–15*) and recent examples of time-cell-like activity during verbal list learning (*16*) and image sequence learning (*17*) tasks, it remains unclear if neurons in humans encode time during task-free conditions that are analogous to those used in animals, or if time cells are restricted to tasks that require explicit attention to time or serial stimuli.

To address these questions, we recorded single- and multi-neuron activity while subjects played a virtual navigation game under strict time constraints. We recruited 10 neurosurgical patients with intracranially-implanted depth electrodes to play a time-constrained, spatial navigation computer game called *Goldmine,* in which they earned points by collecting gold in a visually sparse, underground mine (Fig. 1 and movie S1). Each trial consisted of four, timed events: First, subjects waited passively for 10s at a fixed location, the mine base (Delay_1_). Next, they had 30s to search for gold that appeared throughout the mine in randomized locations on every trial (Gold Search). Subjects then waited an additional 10s in the mine base under identical conditions to the first delay (Delay_2_). Finally, they had 30s to return to remembered gold locations and dig for gold, now invisible (Gold Dig). This sequence repeated for 36 trials per session, and all subjects performed well, collecting 54% (range 35-73%) of golds that were found during Gold Search while maintaining 47% (range 19-71%) digging accuracy (Materials and methods).

**Fig. 1.**
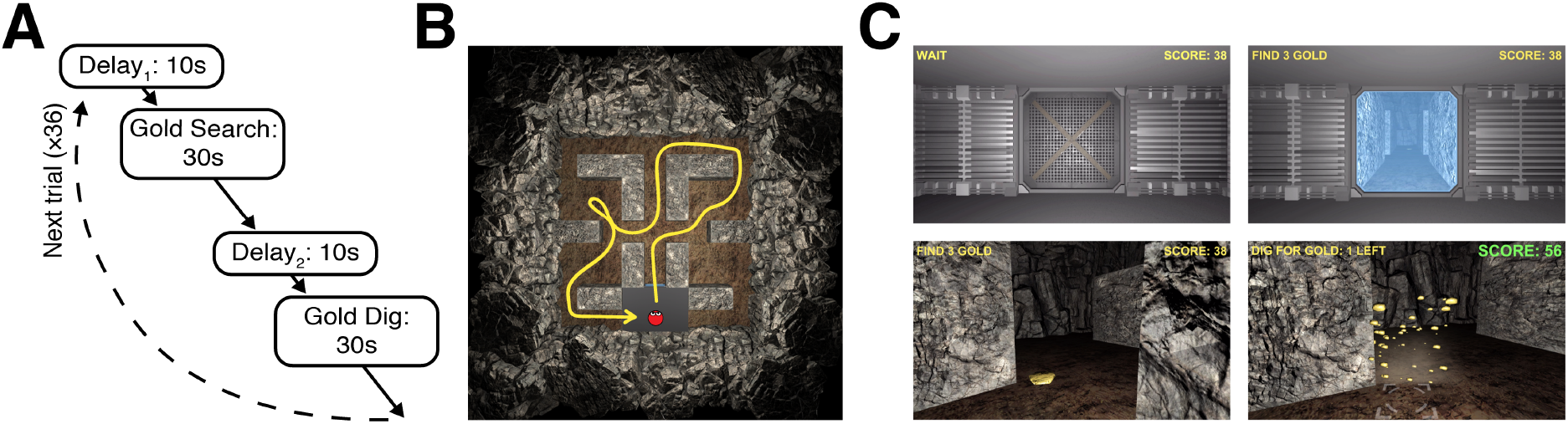
*Goldmine* task. (**A**) Trial structure and timing. (**B**) Top-down view of the mine layout. The yellow line shows an example route by a subject (red circle). (**C**) Gameplay screenshots during Delay (top-left), Gold Search (top-right, bottom-left), and Gold Dig (bottom-right) events.

Using microwires that extended from implanted electrode tips, we recorded extracellular action potentials from 457 single- and multi-units (13-73 units per session, hereafter called ‘neurons’) that were primarily located in the MTL and medial PFC (mPFC; table S1). Here we investigated associations between these neurons’ firing rates and the spatiotemporal structure of the task.

## Time during task-free delays

We first analyzed neural activity during delay events (Fig. 1A), which replicated conditions in which time cells are found in the rodent hippocampus (*9*). To prevent time from being confounded with other behavioral variables, we teleported subjects to the exact same location at the start of every delay, they viewed a static image of a door to the mine that was identical across trials (Fig. 1C), and – as in animal time cell studies – subjects were neither instructed nor incentivized to explicitly attend to time. The delays therefore provided a strict test of the hypothesis that neurons encode time within clearly-defined, repeating intervals with fixed external context.

Figure 2A shows a selection of neurons that fired at precise times consistently across trials, illustrating the range of typical responses. Among the 457 recorded neurons, 99 neurons (22%) exhibited a significant main effect of time (10 discrete, 1s bins), independent of delay-event (Delay_1_ or Delay_2_) and its interactions with time (permutation test against circularly-shifted spikes; see Materials and methods; Fig. 2, A and C). These ‘delay time cells’ were present at rates well above chance in the hippocampus and other recorded regions, with no difference in the proportion of significantly-responding neurons between the hippocampus, surrounding MTL (combining amygdala, entorhinal cortex, and parahippocampal/fusiform gyrus), and mPFC (*P* > 0.05, permutation test controlling for differences in neurons recorded per region, between subjects; table S2). In contrast, only 26 neurons (6%) exhibited a significant time × delay-event interaction, not exceeding chance (Fig. 2C). A majority of time-coding neurons during delays therefore did not distinguish between Delay_1_ and Delay_2_.

**Fig. 2.**
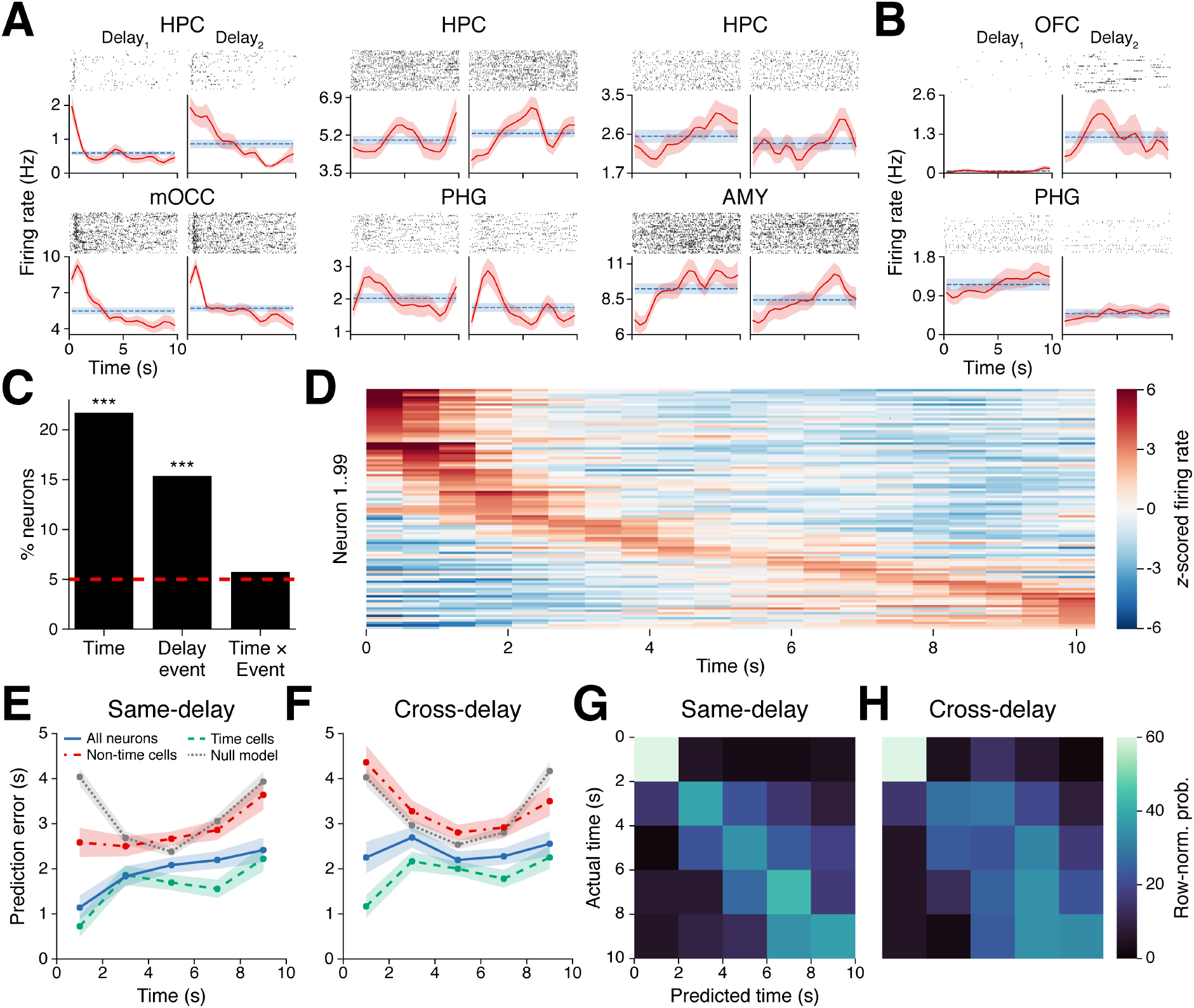
Time cells during task-free delays. (**A**) Subpanels show trial-wise spike rasters and firing rates (mean ± SEM; solid-red line: 500ms moving average; dashed-blue line: grand average) for six time cells in the hippocampus (top row), medial occipital cortex, parahippocampal gyrus, and amygdala (bottom row, L to R). The left subpanel for each neuron shows Delay_1_ activity, and the right subpanel shows Delay_2_ activity. (**B**) Event-specific cells in orbitofrontal cortex (top) and parahippocampal gyrus. (**C**) Percent of neurons with significant responses to each main effect and interaction (red line: Type 1 error rate). *** *P* < 0.0001, binomial test with Bonferroni-Holm correction. (**D**) *z*-scored firing rates for all main effect time cells (each row = one neuron), sorted by time of maximum *z*-scored firing. (**E**) Mean ± SEM prediction errors, across trials, for classifiers that were trained and tested on the same delay event (e.g. both Delay_1_) to decode time from firing rates of all neurons (solid, blue line), the 115 neurons that responded to time as a main effect or interaction with delay-event (dashed, green line), 342 neurons that did not respond significantly to time (dash-dot, red line), and chance-level results from null model classifiers (dotted, gray line). (**F**) Same as (E), but for classifiers that were trained and tested on different delay events (e.g. Delay_1_ → Delay_2_). (**G** and **H**) Confusion matrices for same-delay (G) and cross-delay (H) time cell classifiers. Matrix rows sum to 1, with each value indicating the mean probability across held-out test trials.

Next we examined the distribution of mean firing rates over time for all main-effect time cells, sorted by their maximal firing time (Fig. 2D). Individual neurons had highly variable firing rate peaks, and the number of neurons that peaked in each third of the delay was significantly above chance (*P* < 0.05, binomial tests with Bonferroni-Holm correction). Thus, time cells were not restricted to one portion of the delay but instead spanned the entire 10s duration. However, similarly to time cells in animals (*7, 8, 18, 19*), more time cells peaked near delay onset than at later times (Fig. 2D), and time cells with earlier peaks also showed larger magnitude responses above their baseline activity (*r* = −0.47, *P* < 0.0001; peak firing time versus maximum *z*-scored firing rate).

While time cells did not distinguish between Delay_1_ and Delay_2_, a distinct group of neurons (*n* = 70, 15%) responded to delay-event as a main effect, independent of time (Fig. 2, B and C). Some of these ‘event-specific cells’ had dramatically different firing rates between the two delays, as shown for an orbitofrontal cortex neuron that almost never fired during the 36 Delay_1_ trials yet was active in sustained bursts throughout Delay_2_ (Fig. 2B, top). A similar number of event-specific cells fired more during Delay_1_ than Delay_2_ (*n* = 39, 56%) as showed the opposite response, and like time cells, these neurons appeared throughout the regions we recorded but did not differ significantly between regions (table S2).

Given the prevalence of time cells at the single-neuron level, we next asked if time could be decoded from neural activity patterns at the population level. Indeed, support vector machines trained on firing rates of all recorded neurons decoded time within 1.9 ± 0.1s on held-out test trials, significantly outperforming the 3.2 ± 0.1s error expected by chance (*P* < 0.0001, paired *t*-test versus null model classifiers; see Materials and methods). Mirroring the clustering of time cell peaks near delay onset, classifier error was lowest at delay onset and increased steadily over time, while still remaining better than chance in every time bin (Fig. 2E).

In spatial navigation studies, place decoding is informed both by place cells and ‘non-place’ cells that lack individually-interpretable responses, indicating that spatial location is represented by a distributed neural code (*20*). We compared classifiers that were trained to decode time from all neuron firing rates against classifiers trained only on time cells (*n* = 115, including time as a main effect or interaction) or only on non-time cells (*n* = 342), respectively. Whereas time-cell-only classifiers performed significantly better than all-neuron classifiers (*P* = 0.0070, paired *t*-test; Fig. 2, E and G), non-time cell classifier predictions were no better than chance (*P* > 0.05, paired Ltest). Thus, the population code for time during delays depended on the activity of bona fide time cells, and without these neurons there was no delay time code.

Finally, we considered whether time could be decoded from cross-classifiers that were trained and tested on different delay-events (i.e. Delay_1_ → Delay_2_, or vice versa), as suggested by the prevalence of main-effect time cells over time × delay-event interaction cells. Consistent with this observation, we found that cross-classifiers performed comparably to classifiers that were trained and tested on the same delay event (Fig. 2, F and H). In summary, population neural activity was sufficient to decode time, and the two delays shared an overlapping neural time code.

## Time and place during navigation

Having observed time-coding neurons during delays, we next asked if similar time coding appeared during virtual navigation, when subjects alternated between searching and digging for gold in the mine (Fig. 1A). To identify neural responses to time independent of place, we regressed each neuron’s firing rate against elapsed time (10 discrete, 3s bins), place (12 regions; fig. S1 and S2), navigation-event (Search or Dig), and their first-order interactions. We then identified neurons for which removing the main effect of time or the interaction between time and another variable caused a significant decline in model performance, relative to a null distribution of circularly-shifted firing rates (see Materials and methods). Additionally, to ensure that time and place were sufficiently behaviorally decorrelated, we allowed subjects to exit the base through only one of three doors on a given trial (left, right, or center; counterbalanced across trials), requiring them to vary their routes through the mine. This manipulation severed all but weaker correlations (*r* < 0.2) between temporal and spatial bins, with the exception that subjects always began navigation at the mine base (fig. S3). Models with additional covariates for head direction, movement, visible objects and landmarks, and dig times yielded qualitatively similar results (fig. S4).

Holding place constant, we observed many neurons that fired in a time-dependent manner during navigation (Fig. 3A). Most of these time-coding neurons were context-specific, with a significant number of neurons (*n* = 64, 14%) representing interactions between time and navigation-event, while the number of neurons with a significant main effect of time (*n* = 23, 5%) was at chance level (Fig. 3C). In this respect, time cells during navigation differed markedly from delay time cells that fired analogously between Delay_1_ and Delay_2_. In addition, classifiers trained on neural firing during delays failed to predict time above chance during navigation (fig. S5), and population neural activity was negatively correlated between delay and navigation events, such that neurons that were more active during delays were usually less active during navigation, and vice versa (fig. S6). Insofar as neurons encoded time during navigation, they therefore did not adhere to the delay time code.

**Fig. 3.**
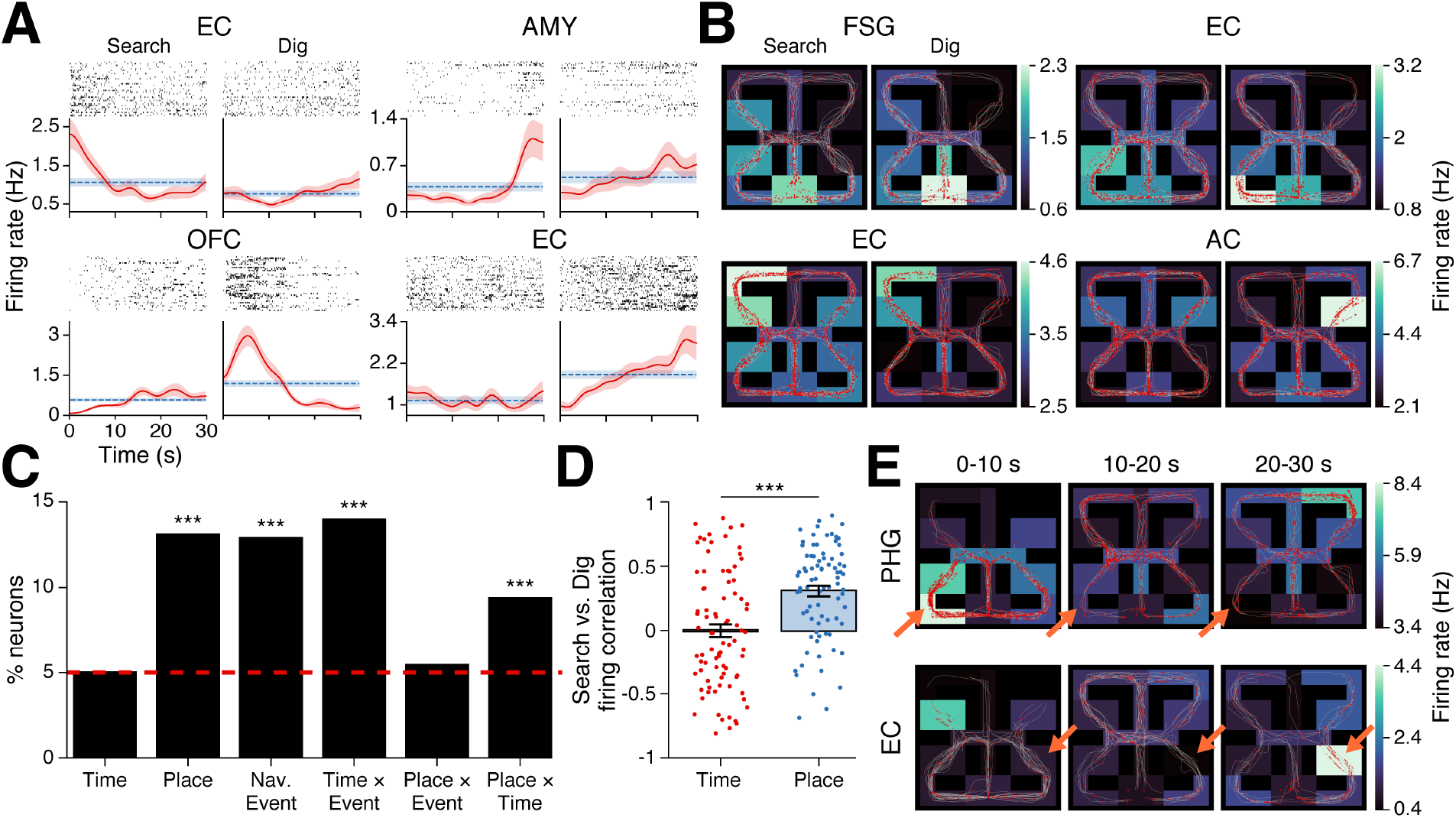
Time and place cells during navigation. (**A**) Four time × navigation-event-specific neurons in the entorhinal cortex (top-left, bottom-right), amygdala (top-right), and orbitofrontal cortex. (**B**) Four place cells in the fusiform gyrus (top-left), entorhinal cortex (top-right, bottomleft), and anterior cingulate. Paths traveled (white lines) and spikes (red circles) are overlaid on firing rate heatmaps (colorbar) in each mine region. The left subpanel for each neuron in (A) and (B) shows activity during Search events, and the right subpanel shows activity during Dig events. (**C**) Percent of neurons that responded significantly to each main effect and interaction (red line: Type 1 error rate). *** *P* < 0.0001, binomial test with Bonferroni-Holm correction. (**D**) Correlated firing rates between Search and Dig events, computed: (Left bar) across time bins, for neurons with a main effect of time or a time × navigation-event interaction (each point = one neuron); (Right bar) across mine regions, for neurons with a main effect of place or a place × navigation-event interaction. *** *P* < 0.0001, Welch’s *t*-test. (**E**) Firing rates in each mine region for two place × time interaction cells in the parahippocampal gyrus (top) and entorhinal cortex, averaged across all Search and Dig events during the first, middle, and last 10s of each event (left to right subpanels).

Most time × navigation-event neurons fired in a temporally-precise manner during one navigation event but had a flat or unrelated firing rate during the other, similar to the place cell phenomenon of ‘global remapping’ (*21*). For example, the entorhinal cortex neuron shown in the bottom-right Fig. 3A subpanel fired at a uniform rate throughout Search events but increased its firing rate more than threefold from the beginning to end of Dig events. As during delays, firing rate peaks for time × navigation-event cells spanned entire event durations and were overrepresented near navigation onset (fig. S7). Time × navigation-event cell proportions did not differ significantly between regions, nor did we find regional differences between other behavioral variables during navigation (table S2).

Classifiers trained on population neural activity during navigation echoed the single-neuron results, decoding time within 3.8 ± 0.2s on held-out test trials (chance: 10.1 ± 0.2s), with increasing error at later times from event onset (fig. S8, A and C). However, contrasting the delay results, time cell cross-classifiers that were trained and tested on different navigation events failed to generalize, performing no better than chance (*P* > 0.05, paired *t*-test; fig. S8, B and D). Thus, while the two delays were represented by an overlapping time cell code, Search and Dig events used orthogonal codes.

As in untimed navigation studies (*13–15, 22*), a significant number of neurons (*n* = 60, 13%) encoded place as a main effect during timed navigation, and these ‘place cells’ exhibited a wide range of receptive fields in different regions of the mine (Fig. 3, B and C). In contrast, the number of neurons with a significant place × navigation-event interaction (*n* = 25, 6%) did not exceed chance. This result revealed a dissociation between time cell and place cell responses to changes in task context, with place cells remaining stable between Search and Dig events while time cells remapped. We confirmed this conclusion by comparing correlations between Search and Dig firing rates: (1) across time bins, for each of the 85 neurons with a main effect of time or a time × navigation-event interaction (*r* = 0.0 ± 0.05 SEM across neurons); and (2) across spatial regions, for each of the 82 neurons with a main effect of place or a place × navigation-event interaction (*r* = 0.31 ± 0.04; Fig. 3D). Despite this remapping dissociation, time cells and place cells did not differ significantly in number or in time and place coding strength, respectively (*P* > 0.05, Welch’s *t*-test comparing likelihood ratio *z*-scores versus null distributions; see Materials and methods).

Prior animal and human studies have found that place cells are often influenced by additional variables including head direction, goal location, and visual cues ^12,21,22^. We asked if time similarly modulates place representations by identifying neurons whose activity reflected interactions between place and time, controlling for their main effects. We identified 43 such place × time cells (9%) whose firing rates at a given location depended on the time it was visited relative to navigation onset (Fig. 3E). These neurons were significantly more prevalent than chance, including in models that further controlled for head direction and visual landmarks (fig. S4). Thus, while time and place were generally represented by different neuronal populations, a small number of neurons conveyed information about joint spatiotemporal context, reflecting a higher level of feature abstraction.

Finally, we identified a significant number of neurons (*n* = 59, 13%) that encoded navigation-event independent of time and place, half of which fired preferentially during Search events and half during Dig events. These navigation-event-specific cells overlapped minimally with delayevent-specific cells described previously (Fig. 2B), and consequently all four trial events were represented by distinct groups of neurons.

## Representing time over long durations

During both delay and navigation events, the neural time code gradually erodes (Fig. 2, E and G and fig. S8, A and C), as reported previously in animals (*18, 23, 24*). Given this loss of temporal information, how do we retain a sense of time over long durations? Behavioral studies of temporal memory suggest that some events might act as ‘landmarks’ in time by realigning the internal clock with the external passage of time (*25*). Landmarks play a parallel role in spatial navigation, where they can correct cumulative path integration error and offer an alternative to direction-based navigation (*26, 27*).

The *Goldmine* task contained two levels of temporal structure: time within each delay and navigation event, and time across the four events in a trial (Fig. 1A). If neurons used the boundaries between these events as temporal landmarks, we reasoned that: (1) it should be possible to decode time across the whole trial at once, and (2) decoding accuracy should decrease steadily within each trial event but increase following the transition from one event to the next.

To evaluate these hypotheses, we trained classifiers to decode time (40, 2s bins) from all neuron firing rates across the 80s trial, then tested them on held-out trials (Fig. 4). As classifiers merely selected the most probable time bin without knowing anything about the trial event structure, above-chance performance can only be explained by the distinctiveness of neural patterns in each time bin, relative to others. Consistent with our first hypothesis, classifier accuracy on held-out trials was well above chance (observed: 25 ± 3% SEM across trials; chance: 2.5%), and classifier-predicted times were closely aligned with actual times throughout the trial duration (Fig. 4B). Consistent with the second hypothesis, classifier accuracy increased sharply at the beginning of each delay and navigation event then decreased at a predictable rate over time (Fig. 4A). Population neural activity was therefore sufficient to decode time across multiple events in the trial.

**Fig. 4.**
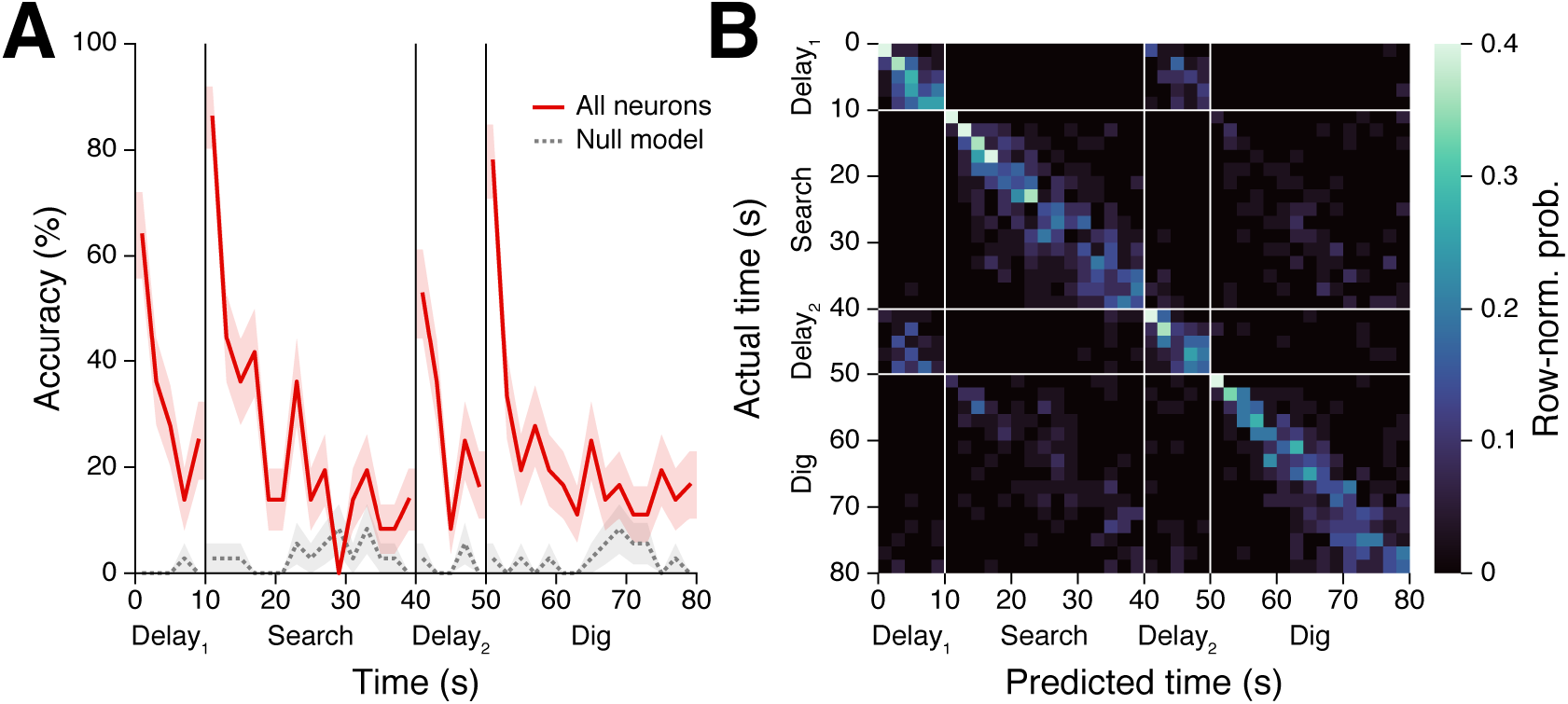
Decoding time across the trial. (**A**) Prediction accuracy by time for classifiers trained to decode 2s time bins from all neuron firing rates, using actual (solid, red line) or circularly-shifted (gray, dotted line) time bins. (**B**) Confusion matrix for classifiers trained on actual (non-shifted) time bins. Matrix rows sum to 1, with each value indicating the mean probability across held-out test trials.

## Discussion

We reveal neurons that encode the passage of time together with space in human brain regions underlying time perception, spatial navigation, and episodic memory. Time cells were active at rest in the absence of movement or other external contextual change, while distinct populations of time cells and place cells emerged during navigation as subjects completed two tasks in a virtual environment for a precise duration.

Fixing the duration of navigation events allowed us to investigate concomitant temporal and spatial codes, and we find the brain maintains largely independent representations of time and place, each encoded by separate populations of neurons within a given context. Our data moreover reveal a novel dissociation between time cell and place cell responses to contextual change, in that most place cells fired similarly between gold searching and digging tasks while time cells completely remapped. This result could reflect differences in how subjects perceived time and place in *Goldmine*, akin to differences in how people judge the time and place of events in daily life. That is, place cells were stable because subjects needed to return to the same locations during gold searching and digging, while time cells remapped to track the temporal progression of these events (first search, then dig) within each trial. This interpretation implies that different experimental conditions could elicit a reversed remapping effect in which place cell activity varies across contexts for which time cells are stable, with potential implications for how events within these contexts are later organized in memory.

Two recent studies in humans described neurons with temporally correlated activity during verbal list learning (*16*) and image sequence learning (*17*) tasks. These studies provided initial evidence for neuronal time coding under conditions in which subjects had to attend to sequential information. Here we extend these findings to show that time codes are present even in the absence of task engagement, serial item presentation, or changing external stimuli. This finding suggests that neurons map time by default, providing a stable scaffold onto which events are bound to their times of occurrence across diverse contexts. Our task additionally enabled direct comparison between human and animal neuronal responses to time, revealing broadly conserved qualities across species. Specifically, we find that human time cells: (1) span entire event durations; (2) accumulate error in the absence of external cues; (3) remap between events for which context discrimination (here gold searching versus digging) is behaviorally adaptive; and (4) reside in the MTL and mPFC (*7–9, 18, 19, 23, 24, 28–31*).

As electrode placement is determined solely by clinical criteria, sampling of different brain regions varied across subjects, and regional differences should be interpreted with caution. In particular, while our results indicate that time cells and place cells are both represented in multiple regions, larger samples are needed to draw conclusions about regional differences. Similarly, caution in interpreting our findings is warranted as subjects are patients with neurological conditions, although consistency with animal studies lends considerable support to these data.

Episodic memory is distinguished from other forms of memory by the recall of events together with the unique, spatiotemporal contexts in which they occurred – the ‘what,’ ‘when,’ and ‘where’ of experience (*10, 11*). Neural representations of these features are thought to converge in the MTL, where neurons fire selectively to image categories and multimodal percepts (‘what’) (*32–34*), and to specific locations and orientations in an environment (‘where’) (*13–15, 22*). Here we confirm a single-neuron basis for knowing ‘when’ an event occurs, separably from knowing ‘where.’ We further show that ensembles of time cells and event-specific cells support a neural code for time across multiple temporal scales. Each element of episodic memory – time, place, and event-related content – is therefore linked to its own neural code, the convergence of which provides a biological foundation for Tulving’s defining view of memory, 50 years ago (*10*).

## Acknowledgments

We are grateful to the participants for their involvement, without which the research would not be possible. We thank members of the Fried and Kahana labs and Marc Howard for helpful discussion and feedback, and we thank Natalie Cherry, Chris Dao, Andreina Hampton, Guldamla Kalender, Connor Keane, and Emily Mankin for assisting in the data collection.

## Author contributions

Conceptualization: DRS

Formal analysis: DRS

Funding acquisition: DRS, MJK, IF

Methodology: DRS

Project administration: MJK, IF

Resources: IF

Software: DRS

Supervision: MJK, ZMA, IF

Visualization: DRS

Writing – original draft: DRS

Writing – review & editing: DRS, ZMA, MJK, IF

## Data and materials availability

All subject-deidentified data are freely available for use upon request to the corresponding authors. Python code and JupyterLab notebooks used for all data preprocessing and analyses can be downloaded at https://github.com/dschonhaut/time_cells.

## Materials and Methods

### Subjects

We analyzed behavioral and single-unit data from 10 neurosurgical patients with drug-resistant epilepsy who completed a total of 12 testing sessions. Clinical teams determined the location and number of implanted electrodes, based on clinical criteria. All testing was performed under informed consent, and experiments were approved by institutional review boards at the University of California, Los Angeles and the University of Pennsylvania.

### Electrophysiological recording

Patients were stereotactically implanted with 7-12 Behnke-Fried electrodes with 40μm diameter microwire extensions (eight high-impedance recording wires and one low-impedance reference wire per depth electrode) that capture local field potentials (LFPs) and extracellular spike waveforms (*35*). Microwire electrophysiology data were amplified and recorded at 30 kHz on a Blackrock Microsystems (Salt Lake City, UT) recording system or at 32 kHz on a Neuralynx (Tucson, AZ) recording system.

### Spike sorting

Automated spike detection and sorting were performed using the WaveClus3 software package in Matlab (*36*). We then manually reviewed each unit for inclusion by evaluating waveform shape, amplitude, and consistency, along with spike time auto-correlation, inter-spike intervals, and firing consistency across the session, and we rejected units that were likely contaminated by artifacts, in keeping with field-standard spike evaluation criteria (*37, 38*). For electrodes with multiple units that passed this inclusion check, we merged units whose waveform features could not be well-separated in principle components space, retaining for analysis a combination of single- and multi-units. Spike sorting was performed by D.R.S., blinded to electrode recording region, and independently reviewed by I.F.

### Task description

Subjects played a first-person, virtual navigation game called *Goldmine,* in which they explored an underground mine while alternating between searching for visible gold and then digging for this gold, now hidden from view, at remembered gold locations. Testing sessions lasted for approximately one hour, and they consisted of a short tutorial sequence followed by 36 experimental trials. Every trial consisted of four timed and two untimed events in the following sequence:

1. Delay_1_ (10s): Subject waits in the mine base. Game controls are turned off, and the subject sees a static image of the center base door. This image is identical across delays, and the subject is in exactly the same location, facing the same direction, on every delay.
2. Gold Search (30s): A 1s beep denotes the start of the Gold Search event, during which the subject may freely search the mine for one or more golds that appear on the ground in randomized locations on every trial. Concurrent with the beep that signals the start of Gold Search, a 1s instruction message, centered onscreen, tells the subject how many golds there are to find. Game controls are also reactivated, and one of three doors (to the left, right, or center) opens to allow the subject to exit the base. If the left or right door opens, an arrow appears onscreen for 1s (overlapping with the instruction message) to indicate which way the subject should turn.
3. Return-to-base_1_ (variable time): All gold vanishes, and a message onscreen instructs the subject to navigate back to the mine base. As soon as the subject re-enters the base, or if they were already in the base at the end of Gold Search, the screen goes black for 2s except for a message that instructs the subject to envision the route they will take during the upcoming Gold Dig event. Across all trials, the median duration for this event was 2s (the minimum duration), and the 75^th^ percentile was 8.9s.
4. Delay_2_ (10s): Subject waits in the mine base. As during Delay_1_, game controls are deactivated, and the subject sees a static image of the center base door. Delay_1_ and Delay_2_ are overtly identical events, differing only by their order within the trial sequence.
5. Gold Dig (30s): A 1s beep denotes the start of the Gold Dig event, during which the subject attempts to return to gold locations from the preceding Gold Search event and dig for gold, now hidden from view. Concurrent with the beep that signals the start of Gold Dig, a 1s instruction message, centered onscreen, tells the subject how many golds there are to dig (equal to the number of golds to find during Gold Search). Game controls are reactivated, and the same door that opened during Gold Search is reopened, allowing the subject to exit the base.
6. Return-to-base_2_ (variable time): Digging is disabled, and a message onscreen instructs the subject to navigate back to the mine base. As soon as the subject reenters the base, or if they were already in the base at the end of Gold Dig, the screen goes black for 2s except for a message that instructs the subject to prepare for the upcoming Gold Search event. Across all trials, the median duration for this event was 2s (the minimum duration), and the 75^th^ percentile was 7.4s.

After 36 trials, a “game over” screen appeared and showed subjects their final score, the number of golds successfully dug, and their digging accuracy. Subjects were aware of the 30s time limits during Gold Search and Gold Dig and of a “short waiting time” between each navigation event, but they were never explicitly instructed to attend to time during the experiment. Instead, they were told that their goal was to maximize their score by digging up as many golds as possible, as accurately as possible. They were also asked to remain focused, still, and silent throughout testing – including during delays – unless they needed to ask the experimenter a clarifying question. Voluntary breaks were programmed after 12 and 24 completed trials. Subjects were also taught to press a ‘manual pause’ button if they needed to pause the game for any reason during testing. We did not analyze trials with manual pauses (1.9% of all trials; min = 0, max = 3 per session).

The game was played from a first-person perspective, with the (invisible) avatar being 2m tall and moving forward at a constant 4m/s. Subjects rotated their view by moving the mouse, moved forward by clicking and holding the left mouse button, and dug (during Gold Dig only) by pressing the spacebar. Releasing the left mouse button caused movement to immediately stop, although head rotation was still possible.

Subjects could retrieve one gold on each of the first two trials. Thereafter, the number of golds, *n_gold_*, varied such that if a subject had successfully retrieved all golds on both of the last two trials, the next trial would have *n_gold_* + 1 golds. However, if the subject failed to retrieve all golds on both of the last two trials, respectively, the next trial would have *max*(*n_gold_* – 1, 1) golds. Otherwise, the number of golds stayed the same. Subjects received 10 points for each gold retrieved, with a correct dig occurring anywhere within 4m of the nearest gold. A crosshairs on the ground in front of the subject indicated where their digging was targeted. Each gold could be retrieved only once, and only golds from the most recent Gold Search event could be retrieved. Subjects were not required to move, or dig, if they did not elect to do so. There was no limit on how many digs could be attempted, but every incorrect dig subtracted 2 points from the overall score. The current score was always visible in the top-right corner of the screen, and current task instructions (how many golds to find or dig) were always visible in the top-left corner of the screen.

Before beginning the main experiment, subjects completed a 10min tutorial with 3 practice trials that taught them the game rules and controls and allowed them to practice moving through the virtual space. The tutorial occurred in a different environment than the one used during the main experiment, although trial events used the same timing (10s delay and 30s navigation events).

The virtual environment was designed to be moderately challenging for patients to learn, capable of being fully explored within 30s, and visually sparse so as to minimize behavioral confounds with time and place. The environment was 27m long × 27m wide and featured 531m^2^ of traversable space, including 477m^2^ in the mine and 54m^2^ in the base. The environment was vertically, horizontally, and diagonally symmetrical except for the base, which served as the sole orienting landmark. Tall rock walls surrounded the mine on all four sides, and inner rock walls were of uniform height (4m) and appearance. The floor of the mine was flat and evenly patterned. Each gold occupied 1m^2^ of space, and all golds were visually identical. Gold locations were selected by the computer at random at the start of every trial, with the only condition being that gold could not overlap with the base, any walls, or any already created golds.

*Goldmine* was programmed in Unity, with scripts written in C#. The paradigm ran on a Macbook Pro at 60 frames-per-second. A cord connected the laptop to a digital-analogue converter that sent patterned pulses to the recording system, for the purpose of later synchronizing electrophysiological and behavioral data.

### Single-neuron responses to task variables

#### Delay events

For each neuron, we used ordinary least squares regression to fit the number spikes (unsmoothed) in 1s increments across the 36 Delay_1_ and 36 Delay_2_ events, as a function of time (10 discrete bins, each 1s in duration), delay-event (Delay_1_ or Delay_2_), and their interaction. We then calculated the likelihood ratio, *LR,* between this model and 3 reduced models that dropped the parameters for each main effect and interaction, respectively. For example, in the case of time, we compared the original model to a reduced model in which all time-bin parameters were removed while delay-event and time-bin × delay-event parameters were retained.

*LR* is calculated as:

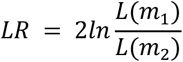

Where *L*(*m_1_*) is the likelihood of the reduced model, given the data, and *L*(*m_2_*) is the likelihood of the original model, given the data. A higher *LR* indicates that the reduced model fit the data increasingly worse than the fit obtained from the original model, and *LR* can therefore be interpreted as a measure of the extent to which a set of parameters (e.g. time bins) improves dependent variable prediction (firing rate) over and above the variance explained by the remaining parameters (delay-event and time-bin × delay-event).

Next, we generated a null distribution for each neuron by shuffling event labels (i.e. permuting Delay_1_ and Delay_2_ labels, without replacement) and circularly-shifting spike counts by a uniform, random integer between 0 and 9, independently across delays. This manipulation served to decouple cross-trial associations between the behavioral parameters and a neuron’s firing rate while preserving both the number of spikes in each time bin and the autocorrelation in firing rates over time. We repeated this process 1,000 times per neuron, recalculating *LR*s between full model and reduced model fits with each iteration. We then compared these null distribution *LR*s to those obtained from the real data, calculating an empirical *P*-value as 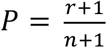, where r is the number of null replicates with an *LR* greater than or equal to the real *LR*, and *n* is the total number of replicates (*39*). We considered a neuron significant for a given main effect or interaction if *P* < 0.05. Finally, we used binomial tests to determine if the number of significant neurons exceeded the 5% Type 1 error rate, Bonferroni-Holm corrected for multiple comparisons across the 2 main effects and 1 interaction term of interest. The results from these models are described in the text and shown in Fig. 2 and table S2.

#### Navigation events

The same procedure was used to analyze firing rate correlations with behavior during navigation as during delays, but with a different combination independent variables. Specifically, ordinary least squares regression was used to model the number of spikes (unsmoothed) in 500ms increments across the 36 Gold Search and 36 Gold Dig events, as a function of time (10 discrete bins, each 3s in duration), place (i.e. subjects’ current location within the 12 mine regions in fig. S1), navigation-event (Gold Search or Gold Dig), and their first-order interactions (time × delay-event, place × delay-event, and time × place). For each neuron, we calculated *LRs* between this model and 6 reduced models that dropped the parameters for each main effect and interaction term, in turn. Empirical *P*-values were obtained relative to null distributions that shuffled navigation event labels and circularly-shifted spike count vectors at random within each navigation event, and neurons were considered significant for a given main effect or interaction if *P* < 0.05. Finally, we used binomial tests to determine if the number of significant neurons exceeded the 5% Type 1 error rate, Bonferroni-Holm corrected for multiple comparisons across the 3 main effects and 3 interaction terms of interest. The results from these models are described in the text and shown in Fig. 3, table S2, and figs. S7 to S9.

We also tested models with additional covariates for virtual head direction (8 angles corresponding to North [the starting direction], Northeast, East, Southeast, South, Southwest, West, and Northwest), player movement (whether the in-game avatar was moving or rotating), base visibility (whether the base was currently visible from the player’s vantage), gold visibility (whether gold was currently visible from the player’s vantage; applied to Gold Search only), gold digging (whether the player had just performed a dig action; applied to Gold Dig only), head-direction × navigation-event, player-movement × navigation-event, and base-visibility × navigation-event. Figure S3 shows the mean correlations, across subjects, between all pairs of behavioral parameters.

### Neural response differences by brain region

As electrode coverage and the number of recorded neurons per region varied between subjects, we used a permutation-based method that accounted for between-subject differences to analyze regional differences in the proportion of significantly-responding neurons to each behavioral variable of interest (see table S2). For each behavioral variable, we first performed a chi-square independence test on the contingency table that listed the number of significantly-responding neurons by region, across all subjects. We then shuffled the neuron-to-region assignment at random within each subject and recalculated the chi-square statistic 1,000 times to obtain a null distribution. Lastly, an empirical *P*-value was obtained that described the extent to which the proportion of significantly-responding neurons by region differed to a greater extent than was observed in the shuffled data (*39*). As no *P*-value passed the significance threshold after adjusting for multiple comparisons, no post-hoc tests were performed. We performed this analysis using 3 regions-of-interest: the hippocampus, surrounding medial temporal lobe (combining amygdala, entorhinal cortex, and parahippocampal gyrus/medial bank of the fusiform gyrus), and medial prefrontal cortex (combining orbitofrontal cortex, anterior cingulate, and supplementary motor area). We excluded from these analyses 46 neurons that were located in more sparsely-sampled neocortical regions due to insufficient sample size.

### Classifying time from population neural activity

We used the scikit-learn library (*40*) to train multi-class, nonlinear (radial basis function) support vector machines to identify discrete, 2s time bins based on population neuron firing rates. We trained separate classifiers on Delay_1_, Delay_2_, Gold Search, and Gold Dig events, respectively (Fig. 2, E to H, and fig. S8), as well as training classifiers across all four of these events combined (Fig. 4).

For each of these conditions, we used the following procedure: First, missing firing rates from the 1.9% of discarded trials (see Task description) were replaced using median imputation. Next, we z-scored firing rates across all time bins and trials in a given analysis, separately for each neuron. Lastly, we trained support vector machines using a nested cross-validation (CV) procedure that split data into train/test/validate folds at the trial level (5 inner folds, 36 outer folds). The inner CV served to identify optimal values for two hyperparameters of the radial basis function kernel: C, which determines the strength of parameter regularization; and γ, which determines the radius of influence for each support vector. For each inner fold, we tested 100 pairs of hyperparameter values, each chosen at random from a continuous, log-uniform distribution between 1*e*^−9^ and 1*e*^9^. The best-performing hyperparameter values were then used to retrain a classifier across the 35 train/validate trials and generate predictions on the held-out test trial. This procedure was repeated over each fold of the outer CV, yielding predictions for each time bin, for all 36 trials.

To evaluate classifier performance, we trained null classifiers that replicated the above procedure, but with time bins being circularly-shifted by a uniform, random integer between 0 and 1 minus the number of time bins, independently on every trial. Paired *t*-tests were used to compare mean prediction errors (absolute value of the difference between actual and classifier-predicted times) across trials for classifiers trained on actual versus circularly-shifted time bins.

Cross-event decoders used the same decoders that were trained separately on each trial event, as described above, but then predicted times from neural firing rates during a different event than the one that was used for training. The following cross-decoders were evaluated (train → test):

1. Delay: Delay_1_ → Delay_2_, Delay_2_ → Delay_1_
2. Navigation: Gold Search → Gold Dig, Gold Dig → Gold Search
3. Delay to navigation: Delay_1_ → Gold Search, Delay_1_ → Gold Dig, Delay_2_ → Gold Search, Delay_2_ → Gold Dig. As the event durations differed, we tested three mappings for each of these combinations: First 10s of navigation, last 10s of navigation, and relative time as a percentage of event duration.

### Code dependencies

Neural firing and behavioral data were analyzed using Python version 3.9.7 and JupyterLab version 3.1.745 (*41*), along with the following, open source Python packages: Matplotlib (version 3.0.3) (*42*), NumPy (version 1.19.1) (*43*), pandas (version 1.1.5) (*44*), SciPy (1.5.2) (*45*), seaborn (version 0.11.1) (*46*), Scikit-learn (version 0.23.2) (*40*), and Statsmodels (version 0.12.1) (*47*).

**Fig. S1.**
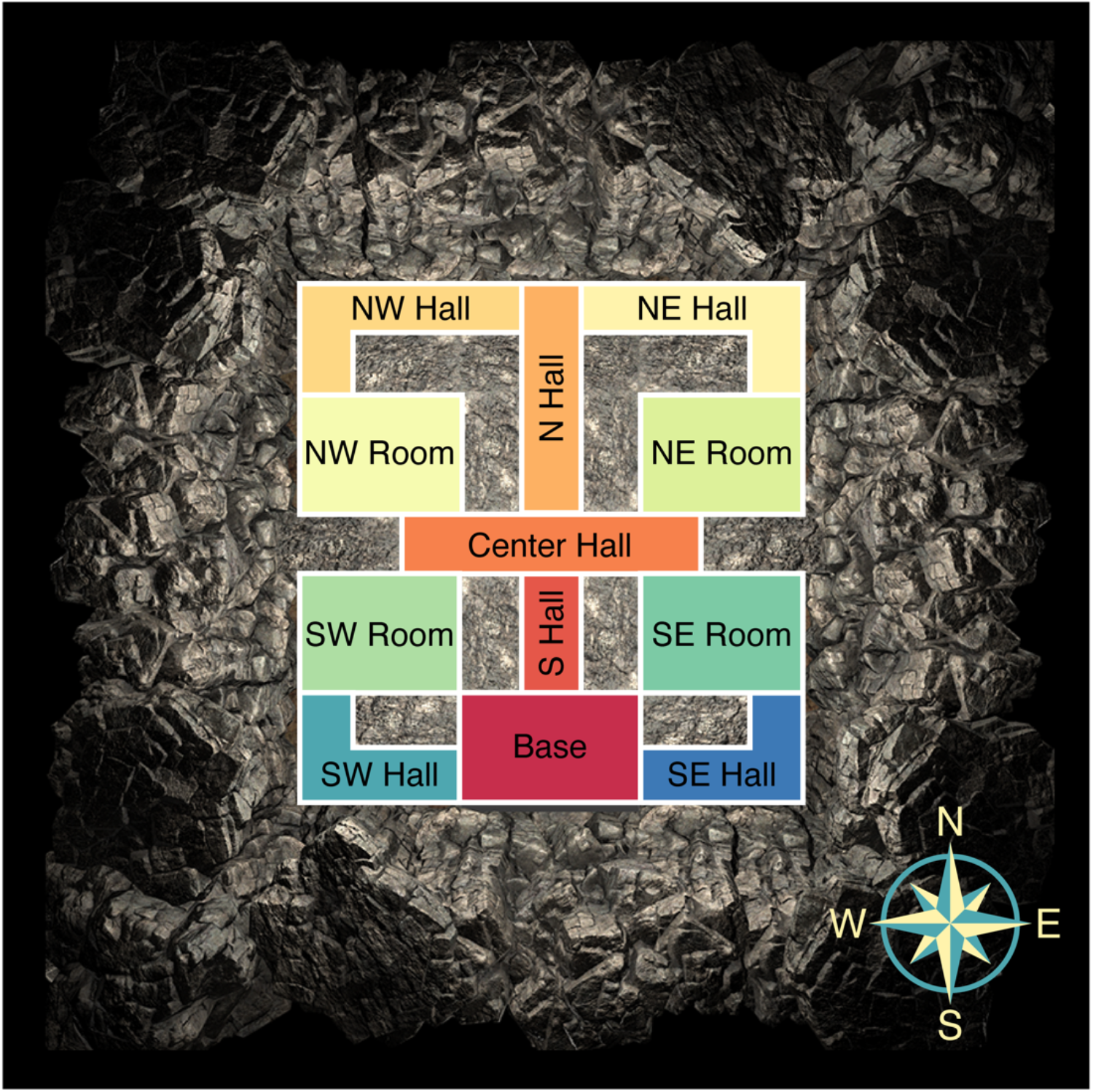
Mine regions. Overhead map of the 12 mine regions that were used in regression models fit to each neuron’s firing rate during Gold Search and Gold Dig events.

**Fig. S2.**
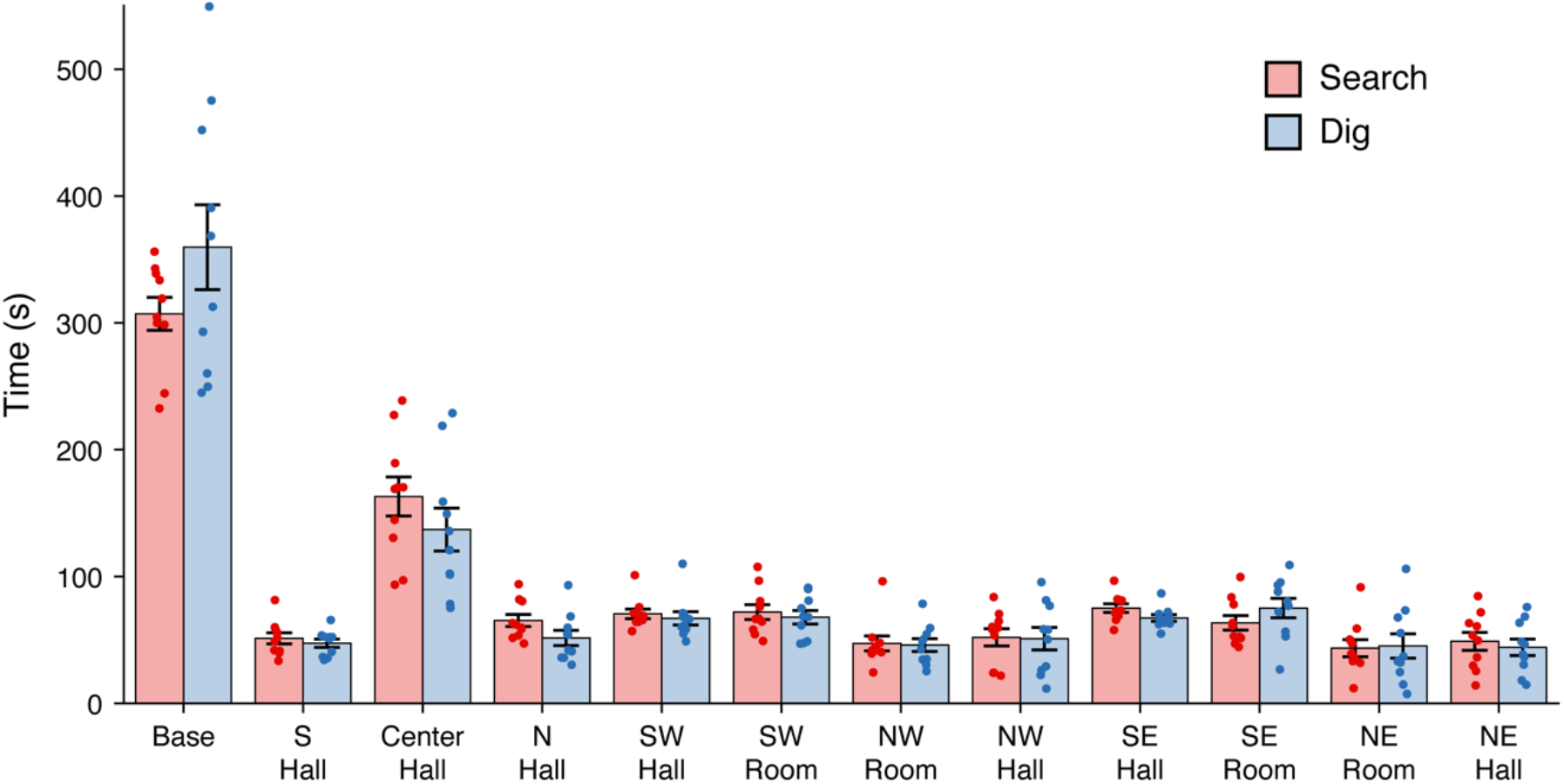
Time spent in each mine region. Bar graph shows how much time subjects spent in each mine region during Gold Search (red, left bars) and Gold Dig (blue, right bars). Bars and error bars show the mean and standard error across subjects, respectively, and overlaid points correspond to individual subjects.

**Fig. S3.**
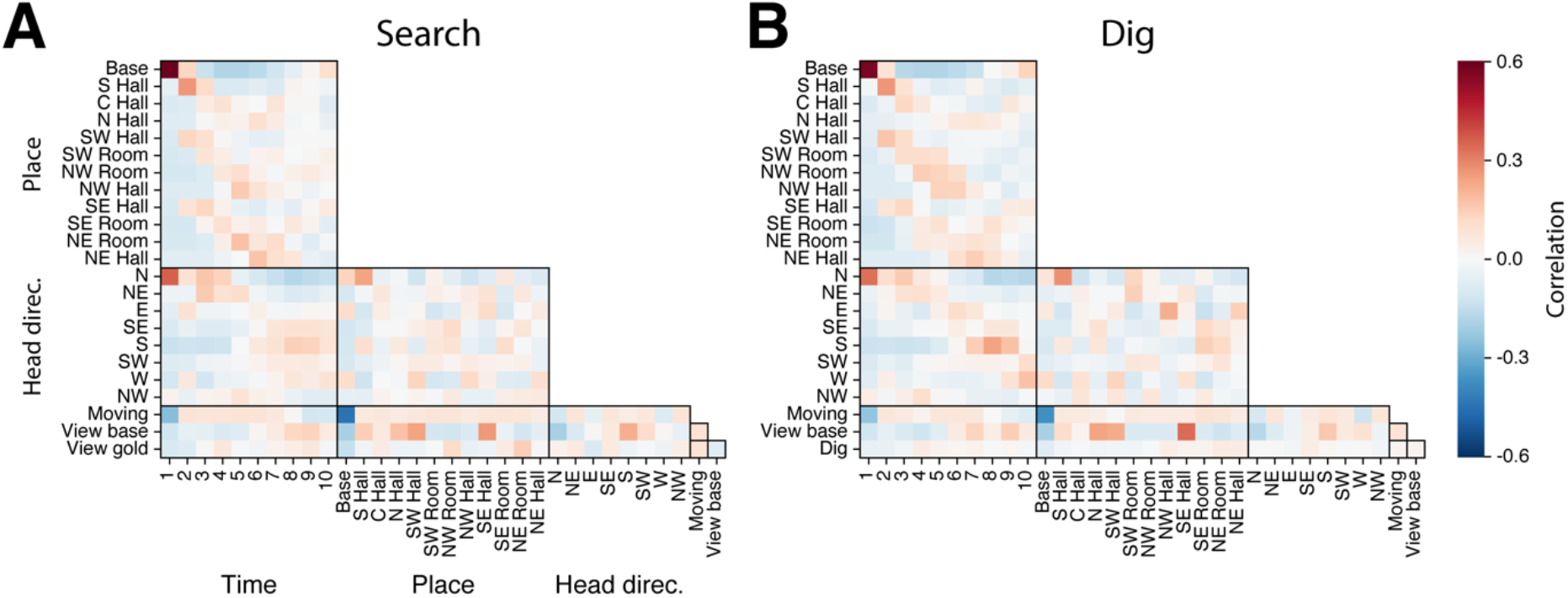
Behavioral parameter correlations. Colormap values show the mean Pearson correlations, across subjects, between each pair of behavioral parameters during Gold Search (A) and Gold Dig (B). Each time bin corresponds to a 3s duration.

**Fig. S4.**
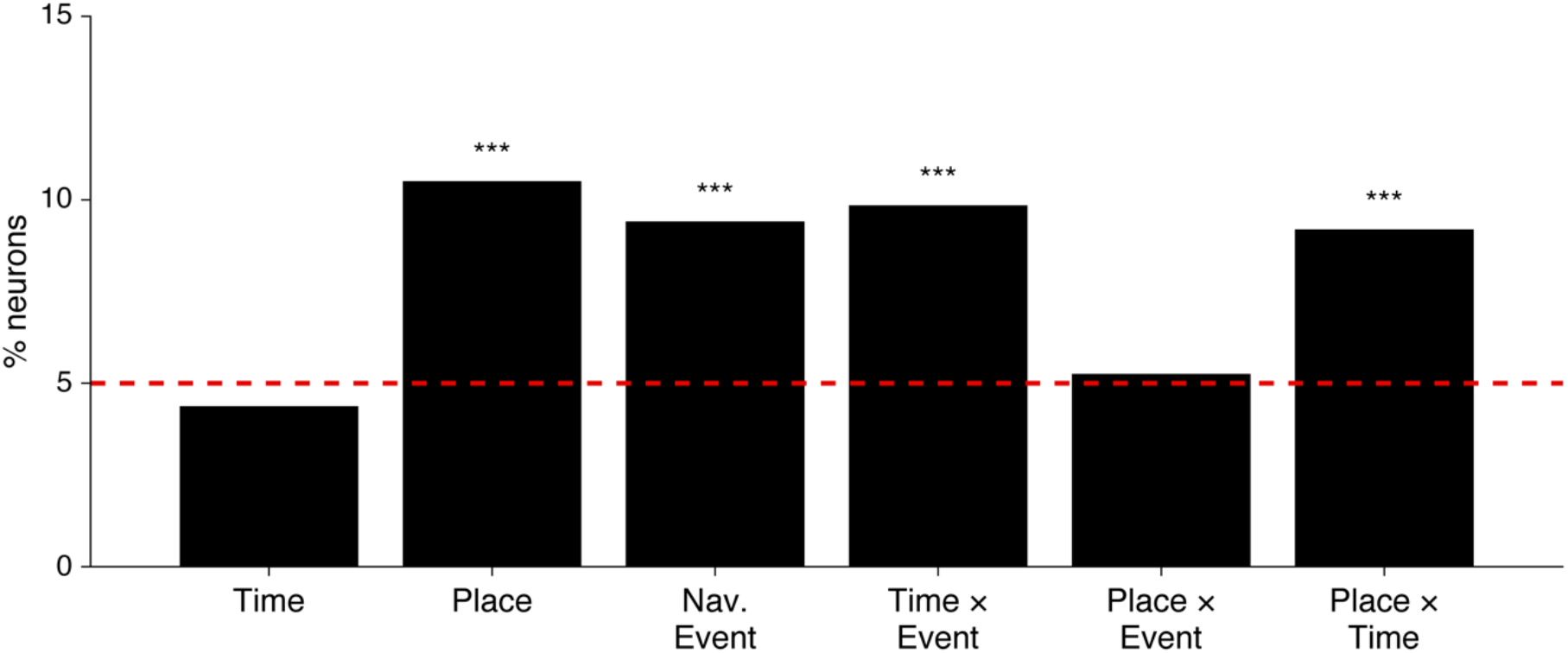
Single-neuron response rates under alternative model specification. Percent of neurons that responded significantly to each main effect and interaction (red line: Type 1 error rate) when adding covariates for head direction, movement, visible objects and landmarks, and dig times. *** *P* < 0.0001, binomial test with Bonferroni-Holm correction. Compare to Fig. 3C.

**Fig. S5.**
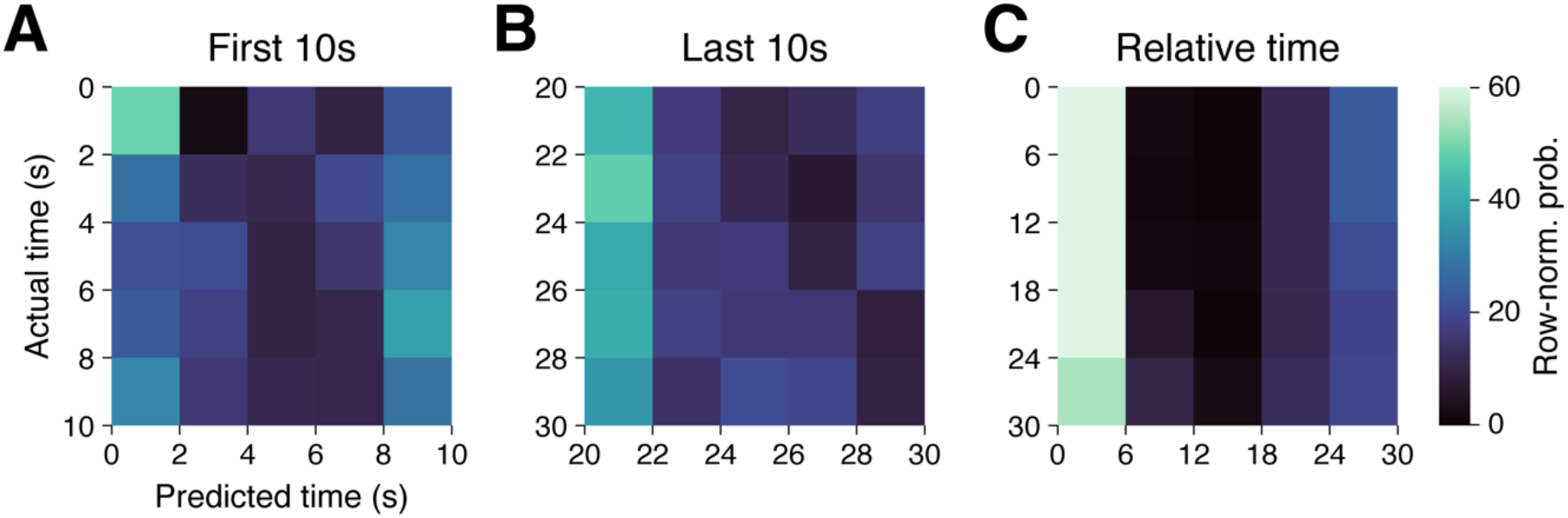
Delay-to-navigation time decoding. Figure panels show confusion matrices for classifiers that were trained to decode time from all neuron firing rates during delay events, then used to predict time during the first 10s (A), last 10s (B), and time relative to event duration (C) during navigation. Matrix rows sum to 1, with each value indicating the mean probability across held-out test trials.

**Fig. S6.**
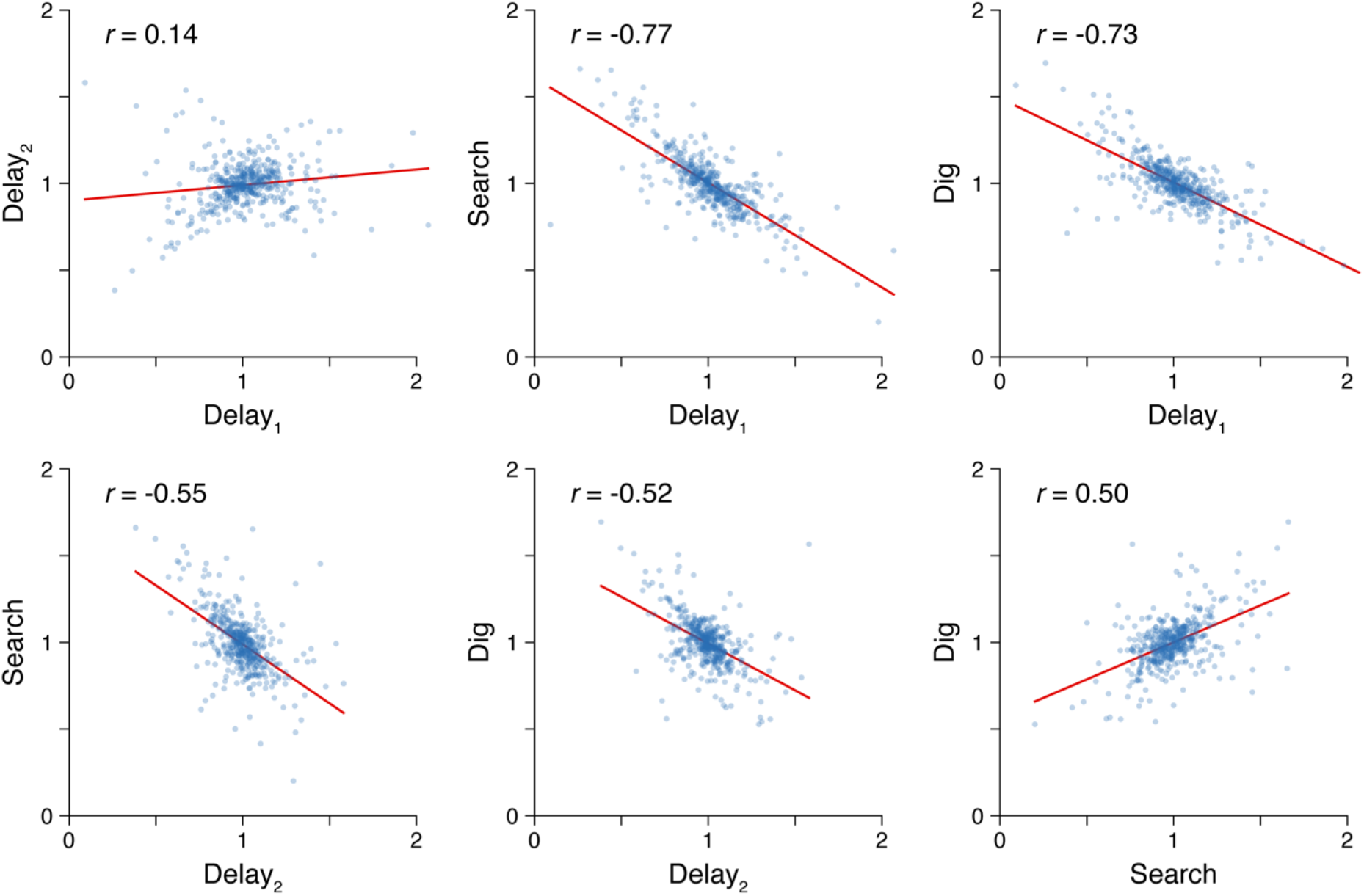
Firing rate correlations between trial event pairs. Figure panels show Pearson correlations between normalized firing rates in each pair of trial events, across all neurons in the study (each dot = one neuron). Each neuron’s firing rate was calculated across time bins and trials within each delay and navigation event, respectively, then normalized by dividing each event’s mean firing rate by the grand average firing rate across the four events.

**Fig. S7.**
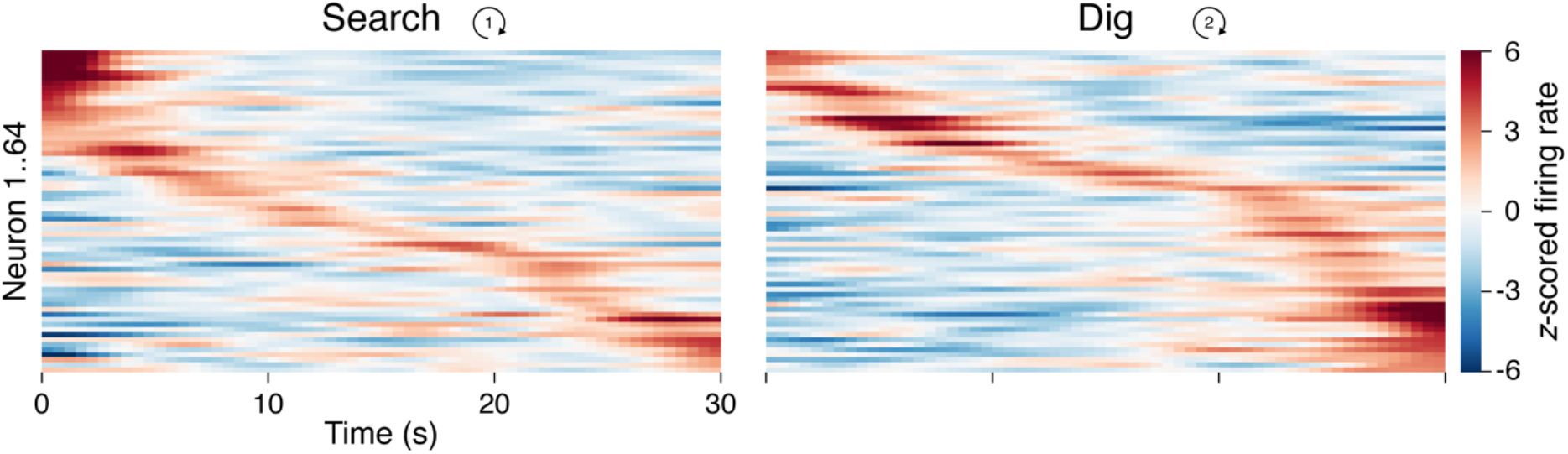
Firing rates over time for all time × navigation-event cells. Heatmaps depict *z*-scored firing rates across the 30s navigation event, averaged across trials, for all time × navigation-event-specific neurons (each row = one neuron), sorted separately by the time of maximum *z*-scored firing during Gold Search (left) and Gold Dig (right).

**Fig. S8.**
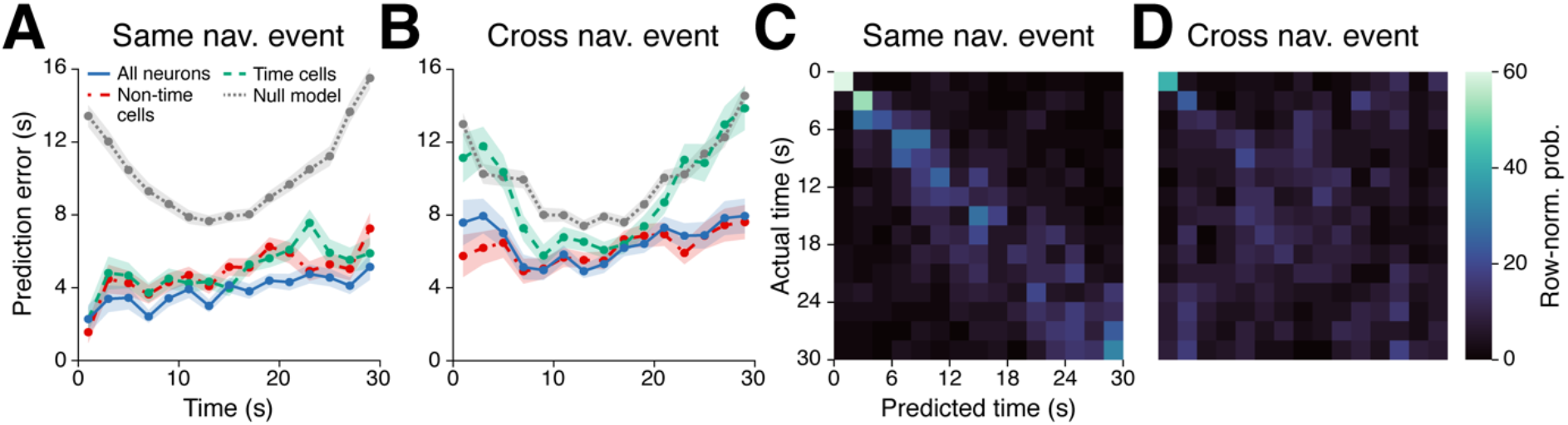
Classifying time during navigation from population neural firing. (**A**) Mean ± SEM prediction errors, across trials, for classifiers that were trained and tested on the same navigation event (e.g. both Gold Search) to decode time from firing rates of all neurons (solid, blue line), the 119 neurons that responded to time as a main effect or interaction with other variables (dashed, green line), 338 neurons that did not respond significantly to time (dash-dot, red line), and chance-level results from null model classifiers (dotted, gray line). (**B**) Same as in (A), but for classifiers that were trained and tested on different navigation events (e.g. Gold Search → Gold Dig). (**C** and **D**) Confusion matrices for same navigation event (C) and cross-event (D) time cell classifiers.

**Table S1.** Neurons by subject and region. Number of neurons (single- and multi-units) recorded from each subject, in each region. HPC = hippocampus; AMY = amygdala; ERCX = entorhinal cortex; PHG/FSG = parahippocampal gyrus/medial bank of the fusiform gyrus; HSG = Heschl’s gyrus; mOCC = medial occipital cortex; mPFC = medial prefrontal cortex (including medial orbitofrontal, anterior cingulate, and pre-supplementary motor area). Subjects are listed in the order tested. * Sample from one patient. ** Subjects with two sessions of data.

| Subject | HPC | AMY | ERCX | PHG/FSG | HSG* | mOCC* | mPFC | Other | Sum |
| --- | --- | --- | --- | --- | --- | --- | --- | --- | --- |
| 1** | 19 | 6 | 5 | 23 | 0 | 13 | 0 | 0 | 66 |
| 2 | 5 | 0 | 0 | 19 | 0 | 0 | 0 | 0 | 24 |
| 3 | 11 | 5 | 1 | 0 | 0 | 0 | 1 | 7 | 25 |
| 4 | 9 | 6 | 10 | 0 | 0 | 0 | 8 | 0 | 33 |
| 5** | 11 | 19 | 42 | 0 | 21 | 0 | 7 | 2 | 102 |
| 6 | 0 | 25 | 24 | 15 | 0 | 0 | 1 | 0 | 65 |
| 7 | 9 | 13 | 15 | 9 | 0 | 0 | 27 | 0 | 73 |
| 8 | 3 | 9 | 0 | 0 | 0 | 0 | 1 | 0 | 13 |
| 9 | 4 | 6 | 3 | 0 | 0 | 0 | 0 | 0 | 13 |
| 10 | 14 | 3 | 0 | 8 | 0 | 0 | 15 | 3 | 43 |
| **Sum** | 85 | 92 | 100 | 74 | 21 | 13 | 60 | 12 | 457 |

**Table S2.** Neuron responses by region. Number and percentage of neurons in each region, across subjects, with significant responses to each behavioral variable. * Sample from one patient.

| **Trial events** | **Behav. variable** | **HPC** | **AMY** | **ERCX** | **PHG/FSG** | **HSG\*** | **mOCC\*** | **mPFC** | **Other** |
| --- | --- | --- | --- | --- | --- | --- | --- | --- | --- |
| Delay<sub>1</sub>,<br>Delay<sub>2</sub> | Time | 10 (12%) | 22 (24%) | 21 (21%) | 22 (30%) | 4 (19%) | 5 (38%) | 13 (22%) | 2 (17%) |
| Delay<sub>1</sub>,<br>Delay<sub>2</sub> | Delay-event | 11 (13%) | 15 (16%) | 25 (25%) | 9 (12%) | 0 (0%) | 2 (15%) | 6 (10%) | 2 (17%) |
| Delay<sub>1</sub>,<br>Delay<sub>2</sub> | Time ×<br>Delay-event | 4 (5%) | 7 (8%) | 4 (4%) | 1 (1%) | 3 (14%) | 2 (15%) | 5 (8%) | 0 (0%) |
| Search,<br>Dig | Time | 2 (2%) | 4 (4%) | 4 (4%) | 8 (11%) | 0 (0%) | 2 (15%) | 2 (3%) | 1 (8%) |
| Search,<br>Dig | Place | 7 (8%) | 6 (7%) | 15 (15%) | 16 (22%) | 1 (5%) | 3 (23%) | 11 (18%) | 1 (8%) |
| Search,<br>Dig | Navigation-event | 8 (9%) | 13 (14%) | 12 (12%) | 12 (16%) | 1 (5%) | 2 (15%) | 10 (17%) | 1 (8%) |
| Search,<br>Dig | Time ×<br>Nav.-event | 9 (11%) | 10 (11%) | 22 (22%) | 8 (11%) | 0 (0%) | 2 (15%) | 12 (20%) | 1 (8%) |
| Search,<br>Dig | Place ×<br>Nav.-event | 4 (5%) | 3 (3%) | 6 (6%) | 4 (5%) | 1 (5%) | 0 (0%) | 6 (10%) | 1 (8%) |
| Search,<br>Dig | Place ×<br>Time | 5 (6%) | 8 (9%) | 8 (8%) | 8 (11%) | 6 (29%) | 4 (31%) | 4 (7%) | 0 (0%) |

**Movie S1.**

***Goldmine* task.** The video shows two complete trials of *Goldmine* as experienced by the test subject.

